# Inbreeding and learning affect fitness and colonization of new host plants, a behavioral innovation in the spider mite *Tetranychus urticae*

**DOI:** 10.1101/2021.06.29.450353

**Authors:** Caroline M. Nieberding, Aubin Kaisin, Bertanne Visser

## Abstract

Habitat fragmentation increases the isolation of natural populations resulting in reduced genetic variability and increased species extinction risk. Behavioral innovation through learning, i.e., the expression of a new learned behavior in a novel context, can help animals colonize new suitable and increasingly fragmented habitats. It has remained unclear, however, how reduced genetic variability affects learning for colonizing more or less suitable habitats. Here, we show that inbreeding in a subsocial invertebrate, the spider mite *Tetranychus urticae*, reduces novel host plant colonization and reproductive fitness. When provided with the possibility to learn from previous experience with a host plant species, outbred mites showed aversive learning ability, but inbred mites did not adapt their behavior. We further found a putative general cost of learning in both inbred and outbred mites. Our results reveal that inbreeding affects the learning component of behavioral innovation for host plant colonization.

## Introduction

Reduction in suitable habitats and increased fragmentation due to human-induced rapid environmental changes poses major challenges for many animals (1). Behavioral innovation, where an individual expresses a new behavior in a novel context (2–4), allows animals to respond to human-induced rapid environmental changes by enabling the use of novel habitats (5,6). Animals can discover and exploit new foods or devise new means of avoiding a threat, as in birds colonizing cities (7,8). Behavioral innovation for novel food sources has also been documented in invasive species (9), such as invasive sparrows that are more willing to approach and consume novel foods compared to non-invasive conspecifics (10). While exploratory behavior clearly plays an important role in generating behavioral innovation (3,11), learning can also lead to the use or renewed application of a behavioral phenotype within novel environments or contexts (7,12). Studying learned responses of animals in the context of habitat selection can thus lead to important insights into the ways by which animals can cope with habitat fragmentation.

Habitat reduction and fragmentation reduces population size and consequently the amount of genetic variation within populations (13–16). Reduced gene flow also leads to mating among kin, inbreeding, inbreeding depression and extinction through persistence of detrimental recessive alleles and/or loss of advantageous heterozygosity (17). Inbreeding depression due to reduced population sizes and genetic diversity has been documented extensively in wild animal and plant populations, as well as in humans (18). Butterflies, birds, and plants with small population sizes indeed often show reduced genetic diversity, lower demographic growth, and increased extinction rates (18). We hypothesized that inbreeding can also affect behavioral innovation, including learning ability to colonize new habitats, because there is strong inter- and intraspecific variation in learning ability across animals, and this is partly due to genetic variation in genes underlying learning ability (19). Polymorphism in the foraging gene (*for*) of *Drosophila*, for example, affects a range of phenotypic traits, including learning (20). Moreover, deleterious mutations impairing learning have been identified in *C. elegans* (21). As learning ability has a genetic basis and displays natural variation, negative effects of inbreeding on learning ability and behavioral innovation may thus contribute to the extinction of populations in isolated habitats.

We focused on host plant switching as a behavioral innovation of low magnitude, meaning that only a minor change to an existing behavior is considered a behavioral innovation (2–4,22,23). Host plant switching occurs in insects during oviposition site selection by adult females when they choose new host plants to lay eggs. In arthropods, female decisions on where to lay eggs have massive consequences for fitness and demography (24,25). This is particularly important for herbivorous insects or arthropods with limited mobility, because the egg-laying site simultaneously serves as the offspring’s food source (26). Oviposition site selection is, therefore, a behavior of key ecological significance in reduced and fragmented habitats, because it determines the ability of arthropods to colonize novel suitable habitats (25–27). Host plant switching is further known to be partly due to learning (28).

Here, we test the hypothesis that inbreeding can affect learning and behavioral innovation through colonization of new host plants. The spider mite *Tetranychus urticae* is an excellent model system to test this hypothesis because host plant species used for oviposition and subsequent larval development are very well known and data on preference-performance relationships are available for different host plant species (29,30). *T. urticae* is also haplo-dipoid, where inbred lines can be set up using, for example, mother-son matings for several generations (e.g. Bitume *et al*. 2013). Furthermore, haplo-diploid species are often more resistant to inbreeding depression due to the purging of deleterious alleles in males. Effects of inbreeding on learning are thus expected to be more pronounced in diploid species. *T. urticae* is further known to modify its oviposition behavior based on previous experience with different host plant species (30). We used a previously developed setup to quantify oviposition after dispersal on a leaf patch across an unsuitable habitat (parafilm bridge)[30–32; Figure 1]. Using this setup, we quantified behavioral innovation in inbred and outbred mites for oviposition site selection after dispersal on two new host plant species, tomato and maize, in addition to the preferred host plant, bean (34–36). Using tomato and maize as novel and less suitable habitats, we quantified colonization success through: *i)* the number of females on a new host plant as an indication of dispersal propensity, i.e., the first step toward colonizing a new suitable habitat; *ii)* the number of eggs on the new host plant species as a measure of colonization success in the new habitat; and *iii)* the total number of eggs laid as an indication of overall reproductive fitness. We expected females to prefer bean for dispersal and egg laying, because this plant species is most palatable and was used to maintain all mites prior to experiments. To specifically test for the effects of inbreeding and learning, two experiments were performed. In the first experiment, we compared the colonization success on new host plants between inbred and outbred mites, where we expected inbreeding to impair colonization success of new host plants. We thus predicted that fewer inbred compared to outbred mites would disperse onto new host plants, and that inbred mites would lay considerably fewer eggs compared to outbred mites. In the second experiment, we tested whether learning, through a previous experience with a new host plant species, would change the propensity of inbred and outbred mites to disperse to and oviposit on a new host plant. We expected learning to modify the number of mites and eggs laid for outbred, but not for inbred mites, because reduced genetic diversity in inbred populations would impair learning ability, and as a consequence behavioral innovation for host plant switching.

**Figure 1:**
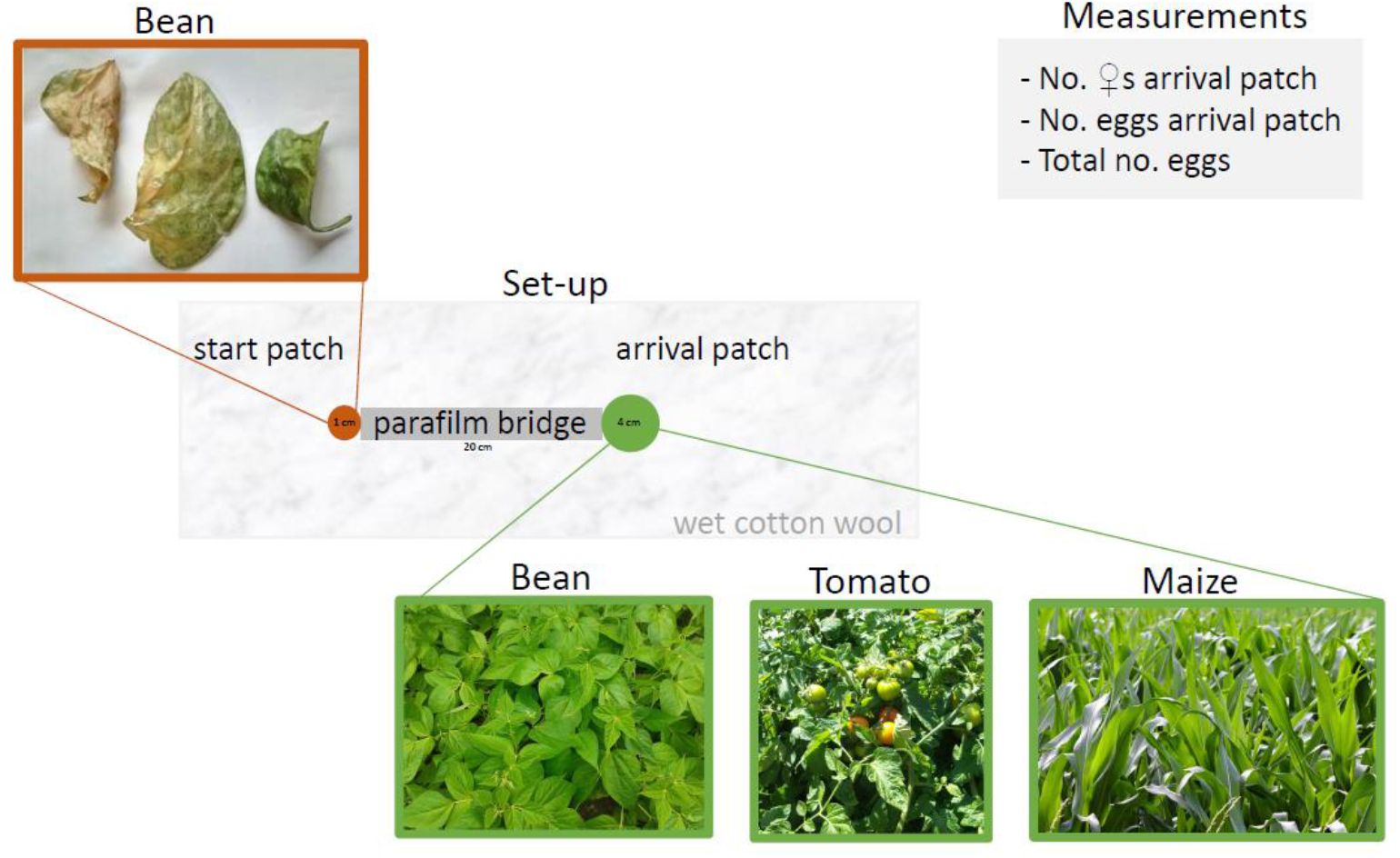
Experimental set-up to test for behavioral innovation through host plant colonization and fitness. Mites were placed on a patch consisting of a degraded bean leaf (“start patch”) that was connected to another patch (“arrival patch”) with the plant species to be colonized (bean, tomato or maize) through dispersal via a parafilm bridge. Colonization and subsequent fitness were determined by monitoring the number of females on the arrival patch (dispersal propensity), the number of eggs on the arrival patch (fitness on the newly colonized plant) and the total number of eggs on the setup (overall reproductive fitness). Mites were tested for their innate and learned colonization and egg-laying behavior.

Our results indeed reveal that inbreeding has a negative effect on host plant colonization by affecting reproductive fitness. We further find that there is a fitness cost to learning. Previous exposure to host plants further led to less colonization by inbred mites and fewer eggs, and oviposition site selection was modified after experience with a new host plant. To the best of our knowledge, these results highlight for the first time that inbreeding interacts with the propensity of behavioral innovation for colonizing new habitats, which has important consequences for predicting how natural populations with reduced genetic diversity will respond to colonize novel suitable habitats in the context of human-induced rapid environmental change.

## Materials and methods

### Model species

*Tetranychus urticae* is a haplo–diploid polyphagous mite that feeds on hundreds of plant and industrial crop species (30,37,38). Depending on environmental conditions, including host plant and temperature, females can lay up to 10 eggs per day for approximately 10 days, a time during which the entire life cycle can be completed. In nature, population density varies from less than 0.1 individual/cm^2^ up to 50 individuals/cm^2^, which depends on the host plant and the state of the infestation (38). Due to a high fecundity and short life cycle, population densities can increase exponentially leading to severe damage or death of host plants. Under controlled conditions, when resources are scarce and population densities become too high, mites disperse individually either by ambulatory or aerial means (39,40). In nature, dispersal of experienced rather than naive animals likely represent the norm, as dispersal can proceed by successive steps where previous exposure to other environments modifies the settlement decision (41,42). Newly emerged 1- to 2-day-old mated adult females are considered to be the dispersers (39) and we previously showed that mated adult females dispersed more when density or genetic relatedness increased on the host plant (32).

An outbred population of *Tetranychus urticae* (strain ‘LS-VL’) was collected from rose plants in a garden near Ghent (Belgium) in October 2000 and maintained on potted bean plants, *Phaseolus vulgaris* (variety ‘Prelude’), with a population size of ∼5000 mites. In addition, multiple inbred lines were produced through a series of mother-son crosses over 20 successive generations in 2012-2013. After 20 generations, these inbred lines were maintained at small population sizes (i.e., 10-30 mated females), leading to a high consanguinity (see Appendix S1 in Bitume *et al*. 2013). Additional inbred lines were further produced by performing mother-son crosses during 1-5 generations in 2018. The inbreeding coefficient for *T. urticae* is expected to be about 0.9 after 4 successive mother-son crosses (29). All inbred lines were maintained in Petri dishes (diameter 10 cm) on bean leaves lying on moist cotton wool. All mites were kept in a climate-controlled room at a temperature of 26.5 ± 1°C, a relative humidity of 60% and a photoperiod of 16:8 L:D.

### Experimental set-up

To test host plant preference of mites, an experimental set-up was designed consisting of a tray with two host plant patches placed on moist cotton wool that acts as a barrier (mites do not move onto that surface), while providing moisture to the leaves. Host plant patches were connected by a 20 cm long piece of parafilm (width = 1 cm; Figure 1). Mites were first placed on a 1 cm^2^ bean leaf disk (“start patch”), and three host plants were used for the other patch (“arrival patch”), tomato, maize or bean (as a control), offered as 4 cm^2^ leaf disks.

### Experimental host plants

Bean, tomato (*Solanum lycopersicum;* Ailsa Craig tomato) and maize (*Zea mays*; variety LG3202) plants were maintained in a phytotron at a temperature of 26.5 ± 1°C, a relative humidity of 60%, and a photoperiod of 16:8 L:D. Leaves of all host plant species used on the arrival patch were collected from the phytotron in good condition, i.e., a state in which the nutritional quality is optimal for a herbivore. In contrast, the bean leaves of the start patch were damaged. Damaged bean leaves were obtained by letting the plants experience prolonged water stress to reduce nutritional quality, and were identifiable through wilting, drying, and yellowing of the leaves. While heavily damaged bean leaves were collected whenever possible, they were not always available due to the need for synchronization between the mites and host plants. The nutritional quality of the bean leaves was recorded before each experiment and considered in the statistical analyses.

### Experimental mites

Mothers at the teleochrysalide stage were collected from inbred lines, as well as the outbred population. Each female was placed on a bean leaf with one male of the same line/population and allowed to lay eggs during 5-6 consecutive days. Each female was transferred to a new bean leaf every 5-6 days (at most 3 times). Offspring were then allowed to develop during the next 10-15 days, after which 1-3-day old adult daughters were collected for experiments. Development of daughters was synchronized to minimize potential age effects.

### Host plant colonization based on innate preference

To test whether females had an innate preference for a host plant species, 10 sisters of the outbred population or inbred lines were placed in the experimental setup (n = 10 replicates for outbred mites, n = 13 replicates for inbred mites). Both density and genetic relatedness can affect dispersal behavior in *T. urticae* (32). We, therefore, aimed to keep density relatively constant at 8-10 individuals/cm^2^, and the coefficient of relatedness kept at ≥ 0.75. The number of females that survived, the number of females on the start and arrival patch, as well as the number of eggs laid on the start and arrival patch was monitored and counted at different times over a ten-day period.

R version 3.6.1 (R Core Team 2020) was used for statistical analyses. Data were analyzed using a Generalized Linear Mixed Model (GLMM) with individual-level variability to account for overdispersion. We analyzed the behavior of females on the setup (number of females on the arrival patch), as well as fitness effects (total egg number and egg number on the arrival patch) with a mixed model and a Poisson distribution. Host plant species of the arrival patch and inbreeding level, as well as their interaction were used as fixed factors in the model, as was female survival in interaction with host plant species of the arrival patch. Replicate nested within family nested within line was used as a random factor, as was the day on which measurements were performed and the degradation level of the start plant. For all statistical analyses, model simplification was used to remove non-significant fixed factors.

### Host plant colonization based on learned preference

To test if females learned to prefer a host plant species during previous exposure, experimental females were obtained and tested as described for the experiment on innate preference above (n = 10 replicates for outbred, n = 19 replicates for inbred mites), with the following exceptions: Females were tested twice using the same host plant species in the arrival patch to allow females to get experience. The ten sisters used in each replicate were maintained on bean plants (control) during 24 hours (experiment day 1), after which they were allowed to move freely between the start and arrival patch during another 24 hours to quantify their naïve host plant preference (step 1 of learning, day 2). Females were then collected and again placed on a fresh bean patch during 24 hours (day 3), after which they were tested a second time during 24 hours (step 2 of learning, day 4). Three out of 10 females were painted with different colors of water paint using a brush, allowing us to track movements of individual females during quantification of innate and learned preferences. On experimental days 2 and 4, patches were monitored after 1, 2, 3, 4 and 24 hours to determine the time to colonization. The number of eggs laid on the setup was counted after 24 hours on day 2 and day 4 of the experiment. We analyzed the behavior of females on the setup (number of females on the arrival patch), as well as fitness effects (total number of eggs and number of eggs on the arrival patch) using a mixed effect model and a Poisson distribution. The level of experience, level of inbreeding and host plant species of the arrival patch were used as fixed factors. Hour nested in replicate nested in family nested in line was used as a random factor, as was the degradation level of the start plant and age of the mites.

## Results

### Inbreeding reduces colonization of new host plants

We quantified colonization of bean, tomato, and maize by counting the number of females and eggs laid after dispersal, as well as the total number of eggs produced (before and after dispersal) as a measure of total reproductive output. Inbred mites colonized bean and maize to a lesser extent than outbred mites, yet more inbred mites were found on tomato compared to outbred mites (Table 1; Figure 2a). Inbreeding reduced the number of eggs after dispersal on the new host plant, and the number of eggs differed between host plant species (Table 1; Figure 2b). There is further a significant reduction in overall reproductive fitness across start and arrival patches (Table 1; Figure 2c), providing evidence that maize is more difficult to colonize than bean. Female survival increased egg laying on bean, but not on maize or tomato (Table 1). Bean, tomato, and maize are thus increasingly more difficult to breed on for *T. urticae* and can be considered a gradient of increasing behavioral innovation for oviposition in relation to habitat selection.

**Table 1:** Inbreeding and host plant species affect colonization propensity and fitness. Innate colonization propensity and fitness were estimated by counting the number of females on the arrival patch, the number of eggs on the arrival patch and the total number of eggs on the set-up (n = 10 replicates for outbred mites; n = 13 replicates for inbred mites). A GLMM was used with line/family/replicate, degradation of the start plant and the day at which measurements were taken as random factors. The relative importance of fixed explanatory variables and their interactions is provided (P ≤ 0.05 in bold).

|  | No females on arrival patch |  | No eggs on arrival patch |  | Total no of eggs (start + arrival) |  |
| --- | --- | --- | --- | --- | --- | --- |
|  | z-value | p-value | z-value | p-value | z-value | p-value |
| Intercept | -0.922 | 0.356 | 2.234 | 0.026 | 3.592 | <0.001 |
| Arrival plant maize | 1.246 | 0.213 | **-2.690** | **0.007** | **-5.751** | **<0.001** |
| Arrival plant tomato | -0.320 | 0.749 | **-1.982** | **0.048** | -1.223 | 0.221 |
| Inbreeding | **1.981** | **0.048** | **2.244** | **0.025** | **2.591** | **0.010** |
| Number of live females | **9.156** | **<0.001** | **7.058** | **<0.001** | **5.903** | **<0.001** |
| Number of live females * inbreeding |  |  |  |  | **-2.926** | **0.003** |
| Arrival plant maize * number of live females |  |  | **-3.493** | **<0.001** |  |  |
| Arrival plant tomato * number of live females |  |  | **-4.875** | **<0.001** |  |  |
| Arrival plant maize * inbreeding | **-2.275** | **0.023** |  |  | 0.594 | 0.553 |
| Arrival plant tomato * inbreeding | -0.806 | 0.420 |  |  | **-2.095** | **0.0362** |

**Figure 2:**
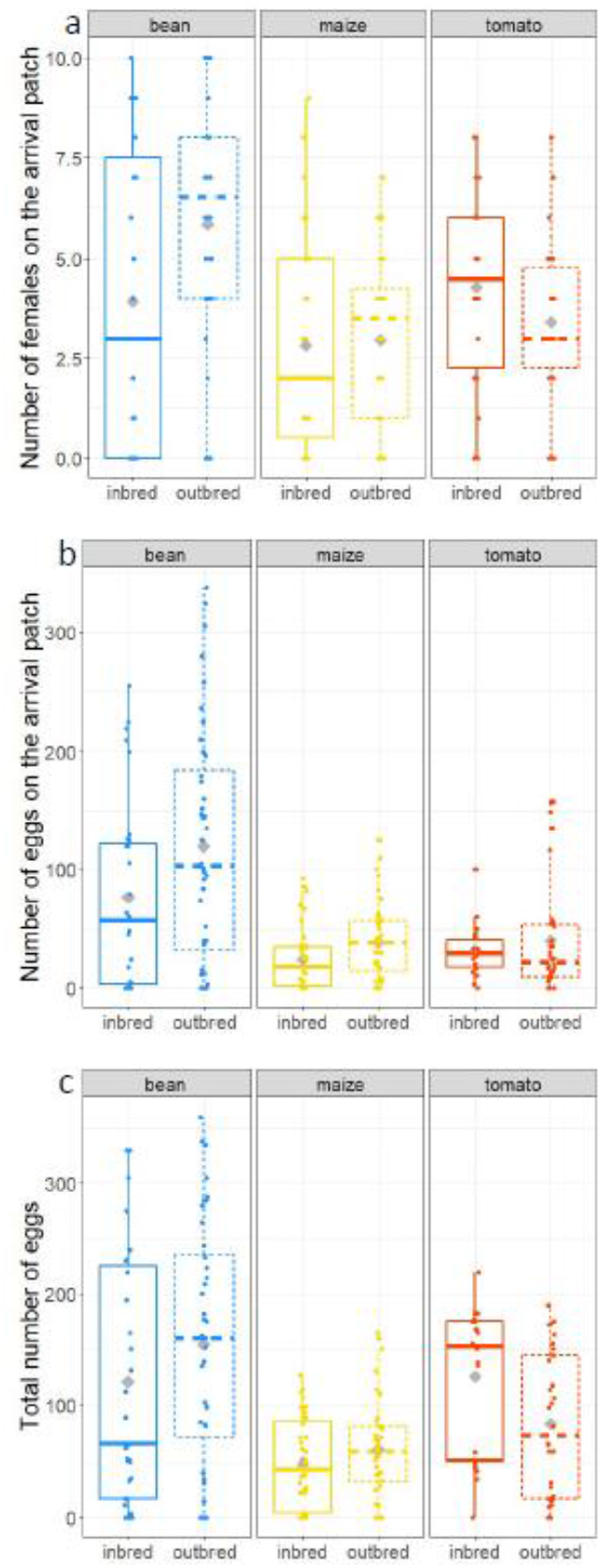
Inbreeding reduces colonization of new host plants. Innate colonization propensity (a) and fitness (eggs on the arrival patch (b) and the total number of eggs (c)) were measured (n = 10 replicates for outbred mites, n = 13 replicates for inbred mites) for inbred (solid lines) and outbred (dotted lines) mites. Inbreeding affected colonization propensity of mites (a), and reproductive fitness was lower on tomato (b) and maize (b,c), meaning that tomato, and to a higher extent maize, are indeed increasingly more difficult to successfully colonize then bean. Statistics are summarized in Table 1.

### Inbreeding affects learning for colonizing new host plant species

Outbred, but not inbred females, dispersed less to a novel host plant after a previous experience with the same host plant, except when colonizing tomato (Table 2; Figure 3a). This suggests that aversive learning took place in outbred mites, and that inbreeding reduced the amplitude of dispersal in response to experience, suggesting reduced learning ability in inbred mites. Experience with a new host plant further led mites to reduce the number of eggs laid on the new host plant (Table 2; Figure 3b). Experience affected inbred mites more than outbred mites, because inbred mites laid fewer eggs compared to outbred mites (with the exception of tomato, where experienced mites -inbred or outbred-laid a similar number of eggs). Learning also affected the total number of eggs laid, with more eggs being produced by naive compared to experienced females (Table 2; Figure 3c), which suggests that our experimental set-up imposed costs on reproductive success for all types of host plant tested. Inbreeding thus differently affects complementary aspects of oviposition site selection: while inbred mites did not learn to modify their dispersal pattern with experience (Figure 3a), their overall reproductive output was modified after experiencing a new host plant (Figure 3b,c).

**Table 2:** Inbreeding and learning interact to affect colonization propensity and fitness. Mites were tested on the same experimental set-up twice to estimate the effect of learning and inbreeding (n = 10 replicates for outbred mites; n = 19 replicates for inbred mites). A GLMM was used with line/family/replicate/hour, degradation of the start plant and age as random factors. The relative importance of fixed explanatory variables and their interactions is provided (P ≤ 0.05 in bold).

|  | No females on arrival patch |  | No eggs on arrival patch |  | Total no of eggs (start + arrival) |  |
| --- | --- | --- | --- | --- | --- | --- |
|  | z-value | p-value | z-value | p-value | z-value | p-value |
| Intercept | -0.691 | 0.490 | 3.810 | <0.001 | 4.891 | <0.001 |
| Step_learning (level of experience) | -0.205 | 0.838 | **-4.198** | **<0.001** | **-5.164** | **<0.001** |
| Arrival plant maize | 0.997 | 0.319 | 0.051 | 0.959 | 0.226 | 0.821 |
| Arrival plant tomato | 1.688 | 0.091 | -0.129 | 0.897 | -1.367 | 0.172 |
| Inbreeding | **2.367** | **0.018** | -0.231 | 0.817 | -0.256 | 0.798 |
| Step_learning * Inbreeding | **-2.490** | **0.013** | **-1.969** | **0.049** | -0.705 | 0.481 |
| Step_learning * Arrival plant maize | -0.805 | 0.421 | **-2.575** | **0.010** | **-4.701** | **<0.001** |
| Step_learning * Arrival plant tomato | -1.180 | 0.238 | **-2.930** | **0.003** | **-3.019** | **0.003** |
| Inbreeding * Arrival plant maize | -0.749 | 0.056 | 0.206 | 0.837 | 0.779 | 0.436 |
| Inbreeding * Arrival plant tomato | -1.911 | 0.158 | -1.381 | 0.167 | -0.240 | 0.810 |
| Step_learning * Inbreeding * Arrival plant maize | 1.412 | 0.158 | -1.663 | 0.096 | **-2.779** | **0.006** |
| Step_learning * Inbreeding * Arrival plant tomato | **2.180** | **0.029** | **4.154** | **<0.001** | **2.360** | **0.018** |

**Figure 3:**
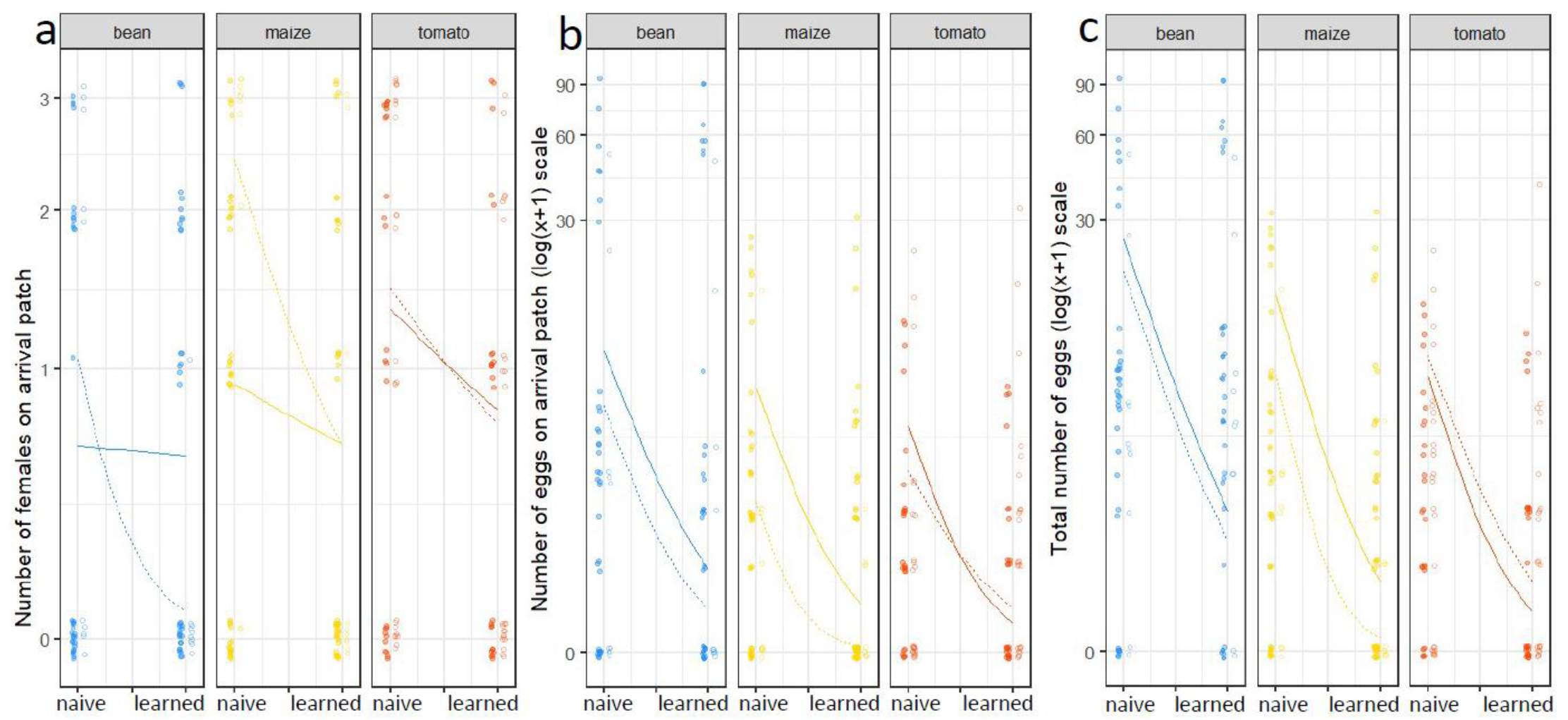
Inbreeding affects learning for colonizing new host plant species. Learned colonization propensity (a) and fitness (eggs on the arrival patch, b, and the total number of eggs, c) were measured (n = 10 replicates for outbred mites, n = 19 replicates for inbred mites) for inbred (solid lines) and outbred (dotted lines) mites. Only outbred females dispersed less to a novel host (a), fewer eggs were laid on the new host plant (b), and learning led to a reduction in the total number of eggs (c). Statistics are summarized in Table 2.

## Discussion

Colonizing new suitable habitats for oviposition and development of the next generation is a behavioral trait of critical importance for herbivorous arthropods. Over the last decades, the coverage of suitable habitats and their connectivity have decreased, reducing the size of, and gene flow among, natural populations (1,13–16). These processes reduce genetic diversity and increase the risk of inbreeding depression. To the best of our knowledge, the effects of inbreeding on learning for oviposition site selection have never been assessed. In this study, we found that fewer inbred female mites colonized a new host plant, and inbred females laid fewer eggs on new host plant species compared to outbred females. This suggests that reduced genetic diversity can severely reduce the chances of colonizing new suitable habitats. We also found that learning, in the form of a first experience with a new host plant, modified the colonization success of a new host plant species differently for outbred and inbred females. Inbreeding indeed reduced learning ability of mites, where a previous experience is an advantage for moving to new habitats. However, when inbred mites did reach the new, harsher habitats, fewer eggs were laid compared to outbred mites. Inbreeding in interaction with learning ability may thus produce a two-fold cost on mites as: *i)* inbreeding increases mortality risks during dispersal because inbred mites do not adapt their emigration propensity to previous experience, and leave anyway; *ii)* inbreeding reduces energy allocation to egg production in newly colonized habitats, compared to experienced outbred mites. The reproductive success of mites is thus affected by complex interacting factors, including experience, inbreeding level, and host plant type.

Are there other mechanistic explanations that could underlie limited dispersal or egg laying on the second experience? The changes in behaviors following experience in our experimental set up cannot be explained by inducible host plant defenses, because for each test we used freshly cut leaf disks. In our experiments, mites of different ages were used and measurements were done at different time points during the day, which could potentially affect motivation and activity level. However, both age and time of experimentation were included as random factors in our statistical models. In addition, females were given 24 hours between learning steps (first and second experience) to rest in order to restore potential reduction in motivation. Moreover, previous work using *T. urticae* revealed that there is little to no variation in egg laying within the range of ages during which our experiments took place (9-14 days; (43)), meaning that there is no evidence that age would reduce egg laying activity or motivation. Mites further form a subsocial species and their dispersal is known to be affected by aggregation and group effects such that all mites within a group do not disperse completely independently from each other (33). In our experiments, the same 10 sister mites were tested for learning ability together throughout the experiment; hence changes in dispersal and oviposition activity were not merely due to an aggregation effect, but due to learning. We show that learning is involved based on the highly significant size effect of the interaction between level of experience (learning steps, i.e., repeated measurements of behaviors between a first and second exposure to a new host plant) and host plant species. Using bean as a control host plant, we found a highly significant change in host plant colonization for the two other plants compared to the effect of bean (Table 2). To the best of our knowledge with regard to the standards for testing learning within the field of cognition (44), we have provided a rigorous test of learning that includes motivation and variation in activity due to age, time of the day, and a control host plant (bean).

What we can further infer from our results is that long term memory (LTM) was involved, which includes protein synthesis (45). Indeed, mites rested for 24 hours before their second exposure, which is the time needed for LTM to build in invertebrates (46). LTM formation is expected to come at a cost, which could explain why fewer eggs were laid during the second exposure for all host plant treatments and inbreeding levels. Costs of learning can also be expressed by delays in reproduction, in addition to reduced fecundity, and both mechanisms could be at play in our experiments (47).

The general question is how inbreeding affects behavioral innovation in relation to colonization of new habitats. The genetic basis of inbreeding depression has largely remained an open question, but one can expect that the expression of overdominant genes, as well as the contribution of many rare, harmful, recessive, and small-effect mutations segregating at low frequencies play a role (17). Learning ability for oviposition site selection is known to show strong inter- and intraspecific variation in animals, and this variation is partly due to genetic variation (19,46). For example, learning for oviposition site selection evolved within a few generations of experimental evolution in the fly *Drosophila melanogaster* (46). A reduced learning capacity in inbred mites could thus be due to increased homozygosity and expression of deleterious recessive alleles of genes responsible for learning ability in oviposition site selection, but this remains to be tested.

In conclusion, our study is the first to show that inbreeding affects behavioral innovation with regard to colonization of suitable habitats and the ability to survive in increasingly reduced and fragmented habitats. While behavioral innovation is traditionally studied in social, long-lived and/or large-brained animals, and behavioral innovation is known to have fitness effects (48), our results can apply to many other species, because short-lived, non-social arthropod species represent the vast majority of all animal species (49). Our results highlight that inbreeding may lead to a negative spiral in the context of habitat reduction and fragmentation, where reduced gene flow, genetic diversity, and inbreeding depression, leads to a reduced ability of inbred females to produce behavioral innovation for reaching new suitable habitats. Inbreeding in natural populations will thus modify colonization ability of new suitable habitats in complex, and so far only poorly understood ways. If we want to be able to quantify the effects of human-induced rapid environmental changes on colonization, we need to better understand how inbreeding, learning and habitat type interact, also in other systems.

## Acknowledgements

We would like to thank Gilles San Martin for his help generating the figures and Christophe Pels for his help with some of the experiments. We are further grateful to Emilie Macke and Isabelle Olivieri for providing inbred lines.

